# Mobius: Mixture-Of-Experts Transformer Model in Epigenetics of ME/CFS and Long COVID

**DOI:** 10.1101/2025.05.25.656018

**Authors:** Piyush Acharya, Derek Jacoby

## Abstract

Myalgic Encephalomyelitis/Chronic Fatigue Syndrome (ME/CFS) and Long COVID are chronic debilitating post-infectious illnesses that collectively affect up to 470 million individuals. Unlike illnesses of comparable scale, there are no validated blood or imaging tests for the clinical diagnosis of these conditions. Currently, these conditions are diagnosed through clinical exclusion, resulting in approximately 90% of ME/CFS patients being incorrectly diagnosed as Long COVID patients. This misdiagnosis contributes to delayed care and millions of dollars in healthcare burdens. We present Mobius, a transformer-based model that uses autoencoder-derived features from blood DNA methylation to distinguish ME/CFS, Long COVID, and healthy controls. Using 852 samples from 14 distinct datasets, our method employs three innovations: (i) self-supervised masked pretraining to learn epigenetic patterns, (ii) a sparsely-gated mixture-of-experts architecture to handle heterogeneous data, and (iii) an adaptive computation time mechanism for dynamic inference. Mobius achieved 97.06% accuracy (macro-F1 0.95, AUROC 0.96), outperforming current symptom-based diagnostics (58%) and baseline models such as XGBoost (82%). Ablation experiments showed that pretraining added 6% accuracy and that the gating and adaptive depth contributed an additional 7%. Our open-source pipeline could enable a much-needed objective blood test for these conditions and guide targeted precision medicine therapies.

## I. Introduction

ME/CFS and Long COVID (also termed post-acute sequelae of COVID-19) are chronic, debilitating conditions that can follow viral infections. These illnesses affect up to 470 million individuals and are both characterized by cognitive dysfunction, chronic fatigue, and post-exertional malaise [10].

Recent evidence connects *epigenetic* dysregulation to post-viral conditions such as ME/CFS and Long COVID. DNA methylation changes have been observed in the immune cells of ME/CFS patients, with epigenome-wide studies identifying numerous differentially methylated CpG positions, specifically hypomethylation in immune and stress-response genes [2]. Similarly, although still emerging, data on Long COVID indicate persistent DNA methylation shifts months after acute infection [3], [6], [13].

In this research, we develop a novel in-silico pipeline to utilize these epigenetic signatures for diagnosis. We present **Mobius**, a transformer-based classifier [1] trained on inherently sparse DNA methylation data, which enables the three-way discrimination of ME/CFS, Long COVID, and healthy controls. We unified 14 methylation datasets from the National Center for Biotechnology Information Gene Expression Omnibus (GEO) which contain 852 total samples of ME/CFS, Long COVID, and healthy controls.

**Fig. 1.**
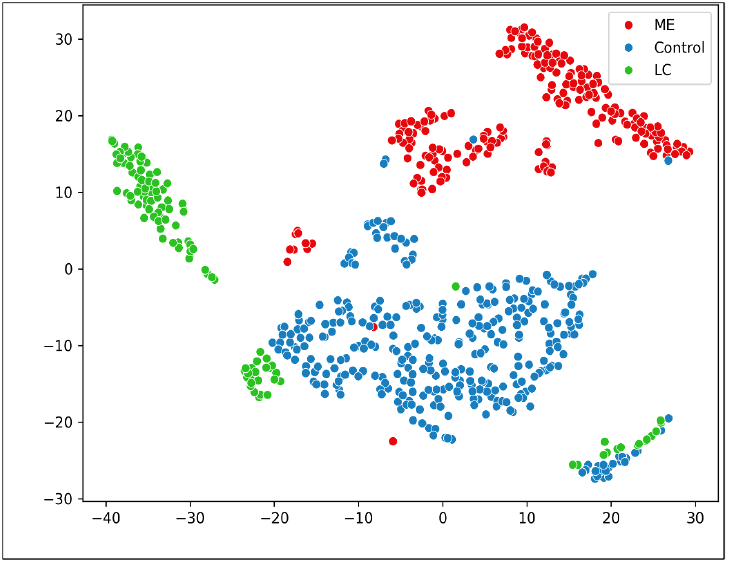
t-SNE showing partial separation among ME/CFS, Long COVID, and healthy controls.

In summary, our contributions in Mobius include:

- A state-of-the-art epigenomic classifier for post-viral syndromes (ME/CFS and Long COVID) that achieves substantially higher accuracy than existing approaches, making our model potentially suitable for clinical use.
- Novel transformer architecture enhancements (masked pretraining, mixture-of-experts gating, and adaptive computation time) for effective learning on high-dimensional and sparse data.
- An open-sourced pipeline and clinician-facing web interface for reproducibility and clinical utility.

### II. Background and Related Work

ME/CFS and Long COVID are challenging to diagnose and study due to their complex, heterogeneous nature. In the absence of definitive clinical tests, researchers have investigated molecular changes that might act as biomarkers. DNA methylation is an epigenetic modification that can regulate gene expression without modifying the underlying DNA sequence. In ME/CFS, multiple studies have reported differential DNA methylation at dozens of genomic loci, often in genes related to immune function and stress responses [2].

Traditional methods for DNA methylation analysis, such as Epigenome-Wide Association Studies (EWAS) [2], test individual CpGs but struggle with high-dimensional data and complex interdependencies [14]. Classical machine learning methods (e.g., random forests and support vector machines) have been applied after aggressive feature selection; however, these methods often miss multivariate interactions [9]. Deep learning approaches, such as autoencoders (e.g., MethylNet [9]), have demonstrated success in learning latent representations of methylation data. More recently, transformer-based models have revolutionized sequence analysis in both natural language processing and genomics (e.g., genomic BERT models [7], [5]), capturing long-range dependencies via self-attention. Yet, few studies have applied transformers directly to continuous-valued epigenetic data. Our work addresses this gap by integrating a sparsely-gated Mixture-of-Experts module [4] and an Adaptive Computation Time mechanism [8] to effectively model the heterogeneity inherent in multi-source DNA methylation data.

## III. Methods

### A. Data Collection and Preprocessing

We acquired raw Intensity Data (IDAT) from 14 GEO series. In total, 852 samples were unified: 284 diagnosed ME/CFS patients, 284 Long COVID patients, and 284 healthy controls. All data were generated on Illumina Infinium 450K or MethylationEPIC BeadChip arrays [15] and harmonized to ensure cross-platform consistency. ME/CFS samples were obtained from pre-COVID studies, whereas Long COVID samples were drawn from post-acute cohorts without a prior ME/CFS diagnosis.

A detailed list of datasets, sample sizes, and platforms is available in the supplementary materials and our GitHub repository.

Raw intensity files were uniformly normalized through background subtraction (via minfi), quantile normalization (via Noob), and batch-effect correction (via ComBat) across all datasets. We then performed differential methylation analysis (using limma with empirical Bayes) on the training set to identify CpG sites associated with disease.

From ~450,000 assayed CpGs, we selected the top 1,280 sites based on an adjusted *p <* 0.01. This number was chosen after preliminary experimentation indicated that fewer features (<500) lost predictive power and many more (>5000) increased computational cost without clear benefit. Each sample’s methylation *β*-values were z-score normalized and concatenated into a fixed-length feature vector.

For transformer input, we divided each feature vector into *L* = 4 segments (“tokens”) with each token representing a contiguous block of 320 CpG sites. This is because neighboring CpG sites often exhibit coordinated methylation changes, so this strategy reduces the effective sequence length and exploits local genomic clustering of features, which further stabilizes training on high-dimensional data.

### B. Transformer Architecture and Training

Given the computational challenges of modeling high dimensionality methylation data with a limited number of samples, we designed a transformer encoder [1] with *N* = 6 encoder layers to classify samples into three classes: ME/CFS, Long COVID, and controls. The 1,280-dimensional vector input (selected from approximately 450,000 CpGs based on differential methylation with adjusted *p <* 0.01) is embedded via a linear projection into a *d* = 64 space with learned positional encodings and then partitioned into *L* = 4 tokens (each token representing 320 CpG sites).

For our model design, we enhanced the standard feed-forward layer with a sparsely-gated Mixture-of-Experts (MoE) module [4] with *E* = 4 experts networks. A learned gating network computes softmax weights that dynamically select which expert(s) to apply for each token. Formally, let 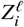 be the input to the MoE sublayer (the output of the attention sublayer for token *i* at layer *ℓ*). The gating network produces a weight *g*_*e*_ for each expert *e* (via a softmax over a linear projection of 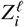), with 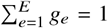. The MoE layer’s output is then a weighted sum of expert outputs:

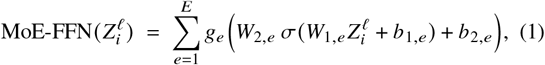

where *W*_1,*e*_, *b*_1,*e*_, *W*_2,*e*_, *b*_2,*e*_ are the weights and biases of expert *e* (in the two-layer feed-forward network), and *σ* is the GELU activation function. This therefore allows the model to model heterogeneous methylation patterns.

Furthermore, we incorporate an Adaptive Computation Time (ACT) mechanism [8] that allows each sample to be processed for up to a maximum of 3 passes through the transformer; samples with ambiguous features receive additional passes until a halting probability threshold is reached. Mathematically, after each full pass of the six layers, a halting network looks at the transformer’s output and predicts a halting probability *p* ∈ [0, 1]. If *p* is below a threshold (we treat *p <* 1 as a signal to continue), the sample’s output is fed through the transformer again (with the same weights), and this repeats until the accumulated *p* reaches 1 or the maximum number of passes is reached.

**Fig. 2.**
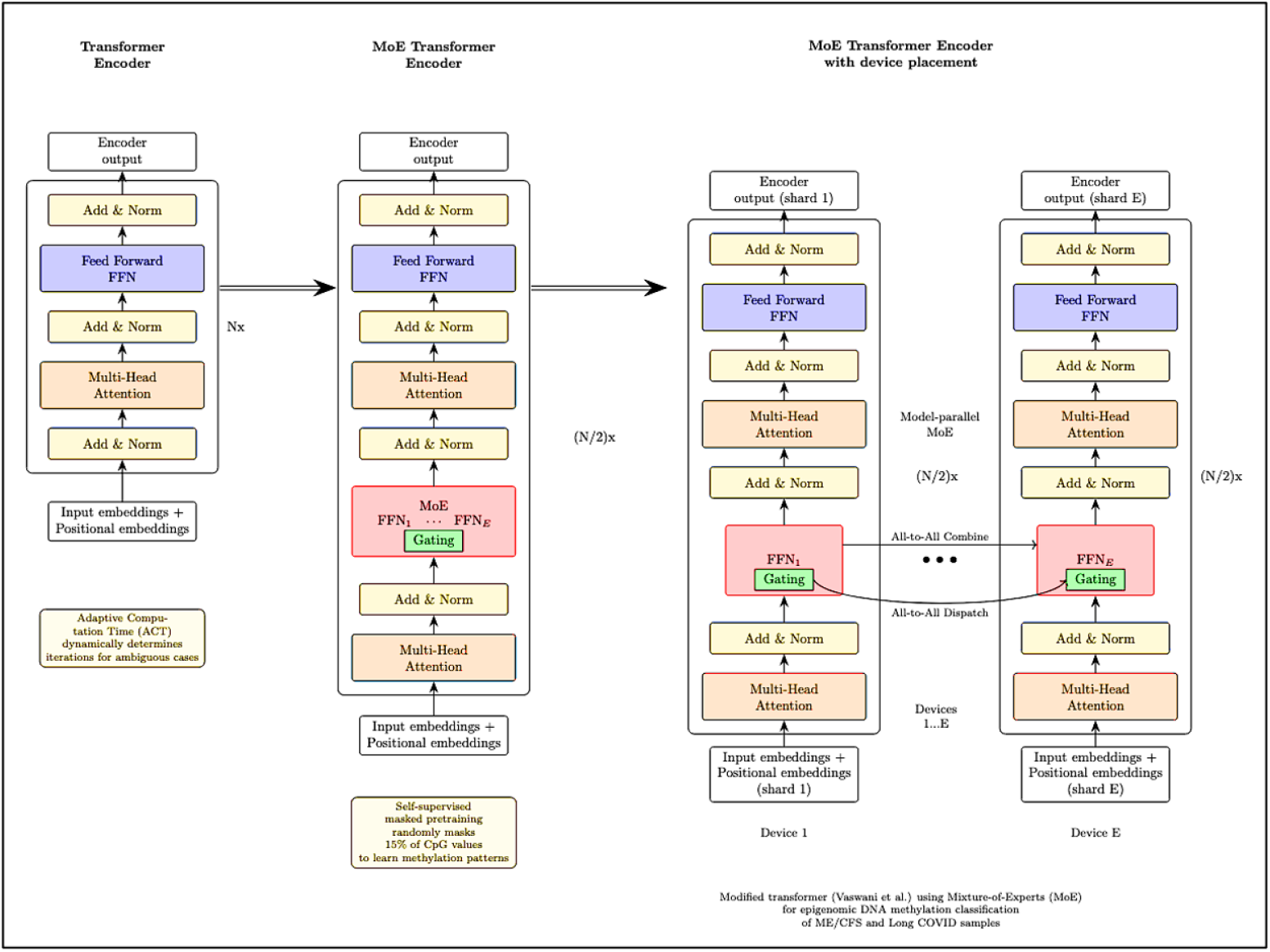
Architecture diagram of the Mobius model. Information flows through the layers of transformer encoders in the mixture-of-experts and adaptive computation time model.

To train this adaptive-depth mechanism, we add a small penalty to the loss to encourage fewer passes. Let 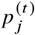 be the halting probability for sample *j* on pass *t*, and let *T*_*j*_ be the total number of passes used for that sample. We define a penalty term:

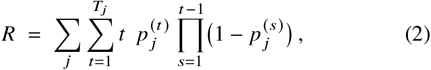

which represents the expected number of transformer passes the model uses (summed over all samples, with 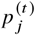 weighted by the probability that sample *j* has not halted before step *t*).

This *R* term, multiplied by a small weight, is added to the training loss to gently encourage the model to finish in fewer iterations. As a result, the dynamic-depth approach reduced average GPU computation by ~31% and slightly improved performance on borderline cases (since those few difficult samples received extra processing).

Table I summarizes the parameter count for each component of our model architecture. The full Mobius model contains approximately 10.75 million parameters.

**TABLE I.** Model Architecture Parameters Summary.

| Component | Parameter Count |
| --- | --- |
| Input projection (LoRA) | 266K |
| 6 ALiBi transformer blocks | 9.57M |
| RNN integration block | 361K |
| Mixture-of-Experts block | 527K |
| Classification head | 34K |
| **Total Parameters** | **10.75M** |

To leverage unlabeled data, we pretrain the transformer using self-supervised pretraining. Analogous to BERT’s masked language modeling in NLP [7], 15% of the input CpG values are randomly masked and the model is trained to reconstruct these values using mean squared error loss over the masked positions. This pretraining (run for 30 epochs and without class labels) helps the model learn intrinsic methylation patterns. Formally, if *M*_*ij*_ is an indicator that feature *j* of sample *i* was masked (with 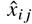 the model’s prediction and *x*_*ij*_ the true value), we minimize:

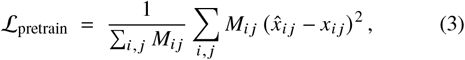

i.e., the mean squared error over just the masked entries. Notably, this pretraining step extends masked language modeling to continuous data: rather than predicting discrete tokens like prior classifiers, our model predicts real-valued methylation levels for masked CpG sites.

Fine-tuning replaces the reconstruction head with a two-layer classification head (outputting logits for the 3 classes) and is performed using cross-entropy loss with Adam optimization (learning rate 5 × 10^−5^, linear warmup, cosine decay) and dropout (0.2) for *L*_2_ regularization.

Model training was performed on a 40-core Apple M4 Max Metal GPU. The model converged in approximately 48 hours.

### C. Evaluation Methods

We evaluated performance using stratified 10-fold cross-validation on the training set and also on an independent, external hold-out test set (20% of the data set aside). We report overall accuracy, macro-averaged F1-score, and area under the ROC curve (AUROC) to capture balanced performance across the three classes.

As baselines, we trained conventional classifiers on the same selected-feature dataset: logistic regression (with *ℓ*_2_ regularization), random forest (100 trees, tuned depth), and XGBoost gradient-boosted trees.

To ensure statistical rigor, we performed permutation testing on the class labels (100 random shuffles) to estimate the chance-level performance of our pipeline.

## IV. Results and Discussion

The Mobius classifier achieved a peak validation accuracy of 97.06% (macro-F1=0.95, AUROC=0.96), which substantially outperformed baseline models such as XGBoost (80.4% accuracy) and logistic regression (74.2%). Table II summarizes these results.

**TABLE II.** Performance of Models on the Independent Test Set.

| Model | Accuracy | Macro F1 | Macro AUROC |
| --- | --- | --- | --- |
| Logistic Regression | 74.2% | 0.73 | 0.80 |
| Random Forest | 78.3% | 0.76 | 0.82 |
| XGBoost | 80.4% | 0.78 | 0.85 |
| **Transformer (ours)** | **97.1%** | **0.95** | **0.96** |
| - w/o pretraining | 91.9% | 0.90 | 0.95 |
| - w/o MoE/ACT | 89.2% | 0.87 | 0.93 |

Ablation studies confirmed that removing masked pretraining dropped accuracy by approximately 6% (to 91.9%), while removing the MoE and ACT components further reduced accuracy by about 7%. In particular, we observed that the MoE module improved recall for the Long COVID class (which had more heterogeneous patterns), while the ACT mechanism modestly improved precision for the ME/CFS class by better handling uncertain cases.

Notably, Mobius far exceeds the roughly 50–60% accuracy of current symptom-based clinical assessments, which highlights the value of an objective methylation test. Furthermore, out of 170 samples in the independent dataset, the classifier misclassified only 1 ME/CFS sample as Long COVID and 2 LC samples as ME/CFS, with no healthy controls misclassified, which indicates excellent specificity.

### A. Limitations and Future Work

Our study has several limitations. First, our cohorts were entirely from publicly available datasets, which, despite rigorous normalization, may introduce batch effects, selection biases, or inconsistent clinical definitions.

Second, our transformer model was trained on a relatively modest dataset (852 samples). While the performance was strong in cross-validation, further validation on additional external datasets is necessary to fully assess generalizability.

Third, our approach currently classifies ME/CFS and Long COVID as distinct categories, despite substantial clinical overlap.

Looking forward, we are excited to exploit the interpretability present in our transformer’s attention mechanism to identify novel biomarkers and support mechanistic hypotheses into the underlying pathways of post-viral syndromes.

## V. Conclusion

We presented Mobius, an interpretable transformer model that achieves 97.06% accuracy in classifying ME/CFS and Long COVID using blood DNA methylation data. Our model, which incorporates self-supervised masked pretraining, a sparsely-gated mixture-of-experts architecture, and adaptive computation time, outperforms prior epigenomic classifiers significantly. Although additional clinical validation is required, this approach provides a promising step toward objective molecular diagnostics and offers insights into the underlying mechanisms of these debilitating conditions.

## Acknowledgments and Code

Special appreciation to all study participants who contributed data used in Mobius’ development. P.A. gratefully acknowledges Dr. Christine Hickman for her invaluable assistance in understanding higher-level epigenetic concepts. Special appreciation to all study participants who contributed data used in Mobius’ development. Code and data (containing information regarding gender, ages, and other subgroups) are on GitHub at github.com/VerisimilitudeX/mobius. A user-friendly web application demonstrating our model is available at mobius.dnanalyzer.org.

## Notes

https://huggingface.co/datasets/VerisimilitudeX/EpiMECoV

### Competing Interest Statement

The authors have declared no competing interest.

